# TIRTL-seq: Deep, quantitative, and affordable paired TCR repertoire sequencing

**DOI:** 10.1101/2024.09.16.613345

**Authors:** Mikhail V. Pogorelyy, Allison M. Kirk, Samir Adhikari, Anastasia A. Minervina, Balaji Sundararaman, Kasi Vegesana, David C. Brice, Zachary B. Scott, SJTRC Study Team, Paul G. Thomas

**Author notes:** these authors contributed equally (Mikhail V. Pogorelyy, Allison M. Kirk). SJTRC Study Team: Joshua Wolf^1^, Aditya Gaur^1^, James M. Hoffman^1^, Tomi Mori^1^, Li Tang^1^, Elaine I. Tuomanen^1^, Hana Hakim^1^, Randall T. Hayden^1^, Diego R. Hijano^1^, Kim J. Allison^1^, E. Kaitlynn Allen^1^, Walid Awad^1^, Resha Bajracharya^1^, Brandi L. Clark^1^, Lee-Ann Van de Velde^1^, Taylor L. Wilson^1^, Ronald H. Dallas^1^, Ashleigh Gowen^1^, Amanda Cole^1^, Jamie Russell-Bell^1^, Ashley Castellaw^1^, Chun-Yang Lin^1^, Maureen A. McGargill^1^, Richard J. Webby^1^, Gang Wu^1^.

## Abstract

ɑ/β T cells are key players in adaptive immunity. The specificity of T cells is determined by the sequences of the hypervariable T cell receptor (TCR) ɑ and β chains. Although bulk TCR sequencing offers a cost-effective approach for in-depth TCR repertoire profiling, it does not provide chain pairings, which are essential for determining T cell specificity. In contrast, single-cell TCR sequencing technologies produce paired chain data, but are limited in throughput to thousands of cells and are cost-prohibitive for cohort-scale studies.

Here, we present **TIRTL-seq (T**hroughput-**I**ntensive **R**apid **T**CR **L**ibrary **seq**uencing), a novel approach that generates ready-to-sequence TCR libraries from live cells in less than 7 hours. The protocol is optimized for use with non-contact liquid handlers in an automation-friendly 384-well plate format. Reaction volume miniaturization reduces library preparation costs to <$0.50 per well.

The core principle of TIRTL-seq is the parallel generation of hundreds of libraries providing multiple biological replicates from a single sample that allows precise inference of both frequencies of individual clones and TCR chain pairings from well-occurrence patterns. We demonstrate scalability of our approach up to 1 million unique paired ɑβTCR clonotypes corresponding to over 30 million T cells per sample at a cost of less than $2000. For a sample of 10 million cells the cost is ∼$200. We benchmarked TIRTL-seq against state-of-the-art 5’RACE bulk TCR-seq and 10x Genomics Chromium technologies on longitudinal samples. We show that TIRTL-seq is able to quantitatively identify expanding and contracting clonotypes between timepoints while providing accurate TCR chain pairings, including distinct temporal dynamics of SARS-CoV-2-specific and EBV-specific CD8+ T cell responses after infection. While clonal expansion was followed by sharp contraction for SARS-CoV-2 specific TCRs, EBV-specific TCRs remained stable once established.

The sequences of both ɑ and β TCR chains are essential for determining T cell specificity. As the field moves towards greater applications in diagnostics and immunotherapy that rely on TCR specificity, we anticipate that our scalable paired TCR sequencing methodology will be instrumental for collecting large paired-chain datasets and ultimately extracting therapeutically relevant information from the TCR repertoire.

## Introduction

ɑ/β T cells are key players in the adaptive immune system, recognizing antigens presented on specialized major histocompatibility complex (MHC) molecules. Both TCRɑ and TCRβ chains are crucial for antigen recognition; the specificity of a TCR is therefore determined by both its TCRɑ and TCRβ sequences, which are encoded by separate transcripts. The TCRɑ and TCRβ chains are formed in a stochastic V(D)J recombination process creating a broad and individually unique ɑβTCR repertoire. T cells recognizing a cognate peptide-MHC complex proliferate during an immune response, forming a clonal population of T cells with the same TCR sequence and antigen specificity. The T cell clone size hierarchy is shaped by antigen exposures and follows a power-law distribution: the largest clone in the repertoire has a frequency of approximately 1-10% of all T cells, while the vast majority of clones are found in just a single cell of a peripheral blood sample. T cell clones expand several thousand fold during the immune response, but the peak frequency of individual clones in most cases is low (around 1 in 10 000, (DeWitt et al. 2015; Pogorelyy et al. 2018) with a combined frequency of all clones recognizing an immunodominant epitope around 1% (Akondy et al. 2009). The great diversity of the TCR repertoire and low frequency of T cells of interest make high-throughput sequencing of TCR genes an essential tool to study T cell responses.

Bulk TCR repertoire sequencing (TCR-seq) technology, introduced over 15 years ago (Robins et al. 2009; Wang et al. 2010), enables the amplification and targeted deep sequencing of TCR repertoires from RNA and DNA, generating datasets with hundreds of thousands of unique T cell clones corresponding to millions of T cells. The major limitation of this method is that it does not provide paired ɑβTCR sequences. Most of the publicly available TCR-seq data is single chain sequencing of TCRβ only, despite both TCRɑ and TCRβ making contacts with the peptide-MHC complex and determining TCR specificity (Garcia and Adams 2005; Krogsgaard and Davis 2005). Although TCRs recognizing certain epitopes might have distinctive features in the TCRɑ or TCRβ chain (Dash et al. 2017; Glanville et al. 2017; Mudd et al. 2022; Meysman et al. 2023), both chains are still needed to confirm specificity *in vitro*.

Concurrently with bulk sequencing, single-cell TCR-seq technologies were developed (Dash et al. 2011; Han et al. 2014; Ludwig et al. 2019) based on single-cell sorting and multiplex amplification of TCR genes in individual wells of PCR plates. Although these methods provide chain pairings and the capability to clone ɑβTCRs of interest for further screening against potential antigens, they are limited in throughput to hundreds of cells, time-consuming, and expensive because of high reagent usage per well.

The development of droplet-based scRNA-seq technologies increased the throughput and lowered per-cell costs for single-cell sequencing (McDaniel et al. 2016; Spindler et al. 2020), with the latest commercially available GEM-X kits from 10x Genomics (Ortolano, n.d.) capable of processing up to 20 000 cells in one reaction. Concurrent reaction miniaturization and transition to a 384-well plate format have reduced the costs of plate-based scRNA-seq (Hagemann-Jensen, Ziegenhain, and Sandberg 2022; Hahaut et al. 2022). Combinatorial barcoding methods (Rosenberg et al. 2018) have further increased the throughput of scRNA-seq up to millions of individual cells. However, high reagent costs and tedious protocols still limit the application of scRNA-seq to a small number of samples. Importantly, most scRNA-seq methods listed above are focused on profiling all polyA transcripts inside a single cell, with optional targeted TCR enrichment.

To improve the throughput of paired ɑβTCR sequencing, Howie et al. suggested a combinatorial approach (pairSEQ) based on even distribution of T cells in a 96-well plate, bulk TCRɑ and TCRβ sequencing of each well, and reconstruction of ɑ/β chain pairings by matching TCRɑ and TCRβ well-occurrence patterns. Lee et al. and Holec et al. suggested alternative data analysis approaches for such pairSEQ experiments. Despite the uniquely high throughput and yield (∼17 million peripheral blood mononuclear cells (PBMCs) were processed in one of the experiments, yielding ∼160 000 unique ɑβTCRs), the protocol was not widely adopted, potentially due to high reagent costs and computationally intensive algorithms for data analysis. Another important limitation for this approach is the inability to pair high-frequency clonotypes, sampled in most or all wells, and low-frequency clonotypes, sampled in only a few wells.

The optimal TCR repertoire sequencing technology should be affordable, able to process millions of T cells (deep), provide complete ɑβTCR sequences allowing for further cloning and screening (paired), and estimate frequencies of individual clones to identify clonal expansion (quantitative). The existing technologies discussed above satisfy only a subset of these requirements.

To address these challenges, we developed TIRTL-seq (Throughput-Intensive Rapid TCR Library sequencing). We combined reaction miniaturization and automation techniques from plate-based scRNA-seq approaches with combinatorial TCR-seq to dramatically reduce the cost and increase the number of T cells analyzed simultaneously. We reimplemented the MAD-HYPE algorithm (Holec et al. 2019) for GPU computation, increasing efficiency and speed by two orders of magnitude, and introduced a novel TCRɑ/β pairing algorithm T-SHELL (TCR ɑβ Sequence Highly Efficient Linkage Learning) which is capable of pairing the most abundant clones in the repertoire. We demonstrate that our method is scalable to ∼1 million unique ɑβTCRs corresponding to over 30 million T cells from a single primary PBMC sample and potentially beyond if more T cells are available.

We validated our results against state-of-the-art bulk and single-cell TCR-seq approaches and show that our method is capable of identifying hundreds of longitudinally contracting or expanding clonotypes and producing paired chains for these clonotypes in a single experiment while also providing biological insight, in this case differential time trajectories for EBV- and SARS-CoV-2-reactive clonotypes.

We believe that wide adoption of our method will lead to explosive growth in the amount of the deep paired ɑβTCR repertoire data acquired, which in turn will accelerate efforts to extract clinically relevant and therapeutically actionable information from TCR repertoires.

## Results

### TIRTL-seq protocol development and optimization

The principle of combinatorial TCR chain pairing is matching TCRɑ and TCRβ occurrence patterns across multiple replicates of T cell samples (Howie et al. 2015; Holec et al. 2019; Lee et al. 2017). This method requires high sensitivity in order to detect transcripts from a single cell within a TCR clone. Furthermore it must be robust enough to load thousands of T cells per replicate and process hundreds of replicates in a cost-effective way.

To achieve these goals, we developed an optimized protocol, TIRTL-seq, to generate bulk TCRɑ and TCRβ libraries from PBMCs or T cells in a 384-well plate format (Fig. 1a). TIRTL-seq utilizes non-contact liquid dispensing with hydrophobic overlays for reaction miniaturization and simultaneous cell lysis and reverse transcription for cDNA synthesis to avoid multiple clean-up steps (Hagemann-Jensen, Ziegenhain, and Sandberg 2022). Targeted amplification of TCRɑ and TCRβ cDNA is carried out in a multiplex reaction using a set of V-segment and C-segment primers from (Howie et al. 2015), which was further optimized to exclude pseudogenes and incorporate parts of Illumina Nextera adapter sequences. The C-segment reverse primers contain 6- or 8-nucleotide long plate-specific barcodes to enable pooling and sequencing of multiple plates in a single sequencing run (see SI Table 1 for oligonucleotide sequences). In the second (indexing) PCR step, we introduce 384-well-specific unique dual indices (UDI) and full length Illumina sequencing adapters. Final libraries are pooled, purified and size-selected using magnetic beads, and quality checked before sequencing on an Illumina platform. The entire protocol takes about seven hours with less than four hours hands-on time from cells to ready-to-sequence libraries and costs about $185 per 384-well plate for reagents and consumables. The default protocol requires a non-contact liquid dispenser and liquid handler to perform 384-to-384 plate transfer and could be implemented on a variety of automation platforms. We also developed a manual version of the protocol using larger volumes in a 96-well PCR plate, which requires no automation and costs about $130/plate (see Methods).

**Figure 1.**
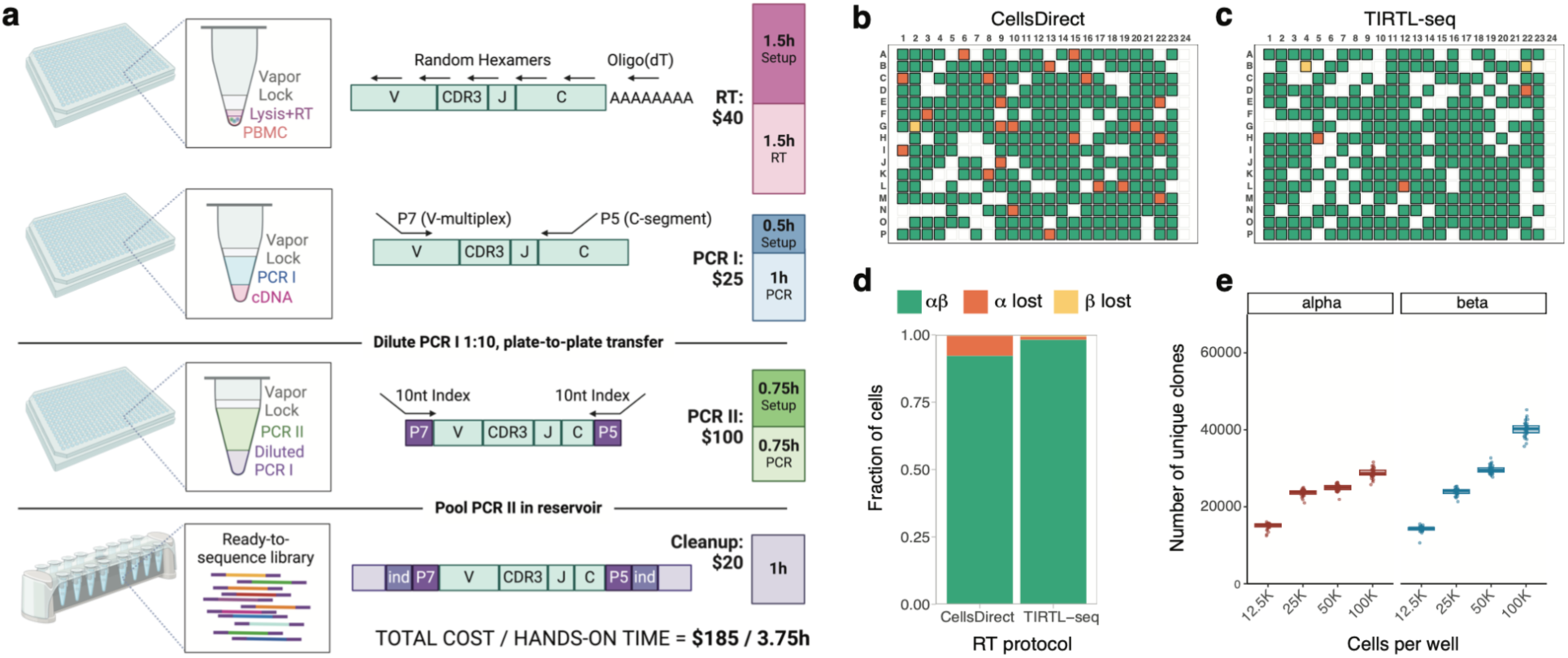
**a.** Schematic of TIRTL-seq protocol. Briefly, a cell suspension is distributed into 384-well plates containing RT/lysis mastermix under a hydrophobic overlay using non-contact liquid dispensers. After the RT reaction, PCR I mastermix with V-segment and C-segment primers is dispensed into the same plate. The PCR I product is then diluted and transferred to the PCR II plate for indexing PCR with well-specific unique dual indices. The PCR II products are pooled by centrifugation, purified, size-selected using magnetic beads, and sequenced on an Illumina platform. Total library preparation cost is listed for one 384-well plate. **b.** CellsDirect vs **c**. TIRTL-seq (Maxima H-based) sensitivity on single sorted T cells. Green: both TCRɑ and TCRβ identified; orange: TCRɑ lost; yellow: TCRβ lost; blank: no cell present. Column 24 is a negative control (no cells sorted). **d.** Relative fraction of cells with both TCRɑ and TCRβ identified (green), lost TCRɑ chain (orange), and lost TCRβ chain (yellow) shown for Invitrogen CellsDirect RT protocol (left) and TIRTL-seq protocol (right). **e.** TIRTL-seq shows robustness to an increasing number of PBMCs. Number of unique clonotypes detected from each well (y-axis) plotted for different numbers of cells per well (x-axis) for the TIRTL-seq protocol.

Next, we tested the sensitivity of TIRTL-seq for detecting TCRɑ and TCRβ transcripts using single-sorted live CD3+ cells from healthy donor PBMCs. As efficient simultaneous cell lysis and reverse transcription (RT) is a crucial step of the protocol, we compared the commercially available proprietary Superscript IV-based CellsDirect kit to our Triton-X100/Maxima H-based cell lysis/RT reaction with the same amplification conditions for PCR I and PCR II. Using the TIRTL-seq RT protocol, we detected both TCRɑ and TCRβ transcripts in over 98% of wells containing single T cells, outperforming the commercially available CellsDirect kit at a fraction of the price (Figure 1b-d). Next, we loaded 12 500 - 100 000 PBMCs per well of a 384-well plate to test the robustness of TIRTL-seq to increasing cell number. We find that the number of unique clonotypes identified increased monotonically with increasing numbers of cells for TCRβ (Fig. 1e). However, the number of unique clonotypes identified plateaued at 25 000 cells/well for TCRɑ, potentially due to the lower number of transcripts per cell (Fig. 1e) (Genolet et al. 2023; Ma et al. 2018).

Overall, the TIRTL-seq protocol proved robust and sensitive across a range of cell numbers up to 25 000 PBMCs per well for TCRɑ and TCRβ. Thus, the maximum number of cells that can be analyzed in a 384-well plate corresponds to approximately 10 million PBMCs, well within the expected counts from a regular blood draw of 1 to 10 mL.

### TIRTL-seq identifies large number of ɑβTCR pairs

To predict ɑβTCR chain pairings in the TIRTL-seq data, we applied the MAD-HYPE algorithm developed by Holec et al. based on the principle of combinatorial pairing by mapping presence/absence patterns originally proposed by Howie et al. Briefly, for each possible TCRɑ/TCRβ pair, the algorithm calculates the number of wells in a plate where chains are detected simultaneously, the number of wells where each chain is found separately, and then accepts or rejects the pair using a Bayesian statistical model. The MAD-HYPE algorithm was originally implemented in Python 2.7, which is no longer supported. We reimplemented the algorithm in R and Python 3 and introduced several critical performance improvements (see Methods). In addition we used the cupy package (Okuta et al. 2017) to run on NVIDIA GPUs and the MLX framework (Hannun et al. 2023) to run on the Apple Silicon platform. Our implementation of the MAD-HYPE algorithm achieved a runtime of 93 seconds for data from Experiment 2 in ref. (Howie et al. 2015) when run on a single NVIDIA GPU A100 80GB in comparison to 24 hours reported for the same dataset in ref. (Holec et al. 2019) on a 96 core CPU cluster, resulting in an almost 1000x increase in computation speed. Using this pairing approach, we identified 169 423 unique ɑβTCR pairs from TIRTL-seq data generated in a 384-well plate loaded with 10 million PBMCs. Using the manual 96-well protocol, we identified 88 142 paired TCRs from 10 million PBMCs per plate.

We then investigated how increasing the number of 384-well plates, each loaded with 10 million PBMCs, prepared from the same donor would increase the number of identified ɑβTCR pairings. As we added additional plates, we tried three alternative strategies to analyze the data: (i) calling pairs independently for each 384-well plate and combining results (“more plates”); (ii) combining wells from multiple plates into a single 384-well plate before analysis to simulate more cells per well ("more cells per well"), and (iii) analyzing all wells collectively ("larger plate with more wells") (Fig. 2a). We found the last strategy to be the most effective in identifying ɑ/β chain pairings. Increasing the number of wells analyzed together improved the resolution of matching ɑ/β chain patterns, resulting in higher statistical power of the approach. We identified 989 241 paired ɑβTCR clonotypes corresponding to more than 30 million cells from six 384-well plates each loaded with 10 million PBMCs. Thus we show that increasing the number of wells per sample increases the number of identified ɑβTCR pairs. On the other hand, decreasing the number of wells per sample in a TIRTL-seq experiment would accommodate T cells from multiple donors in the same plate to further reduce cost for cohort studies. We tested this scenario by downsampling the number of wells. Limiting analysis to 96 wells decreased the number of identified ɑβTCRs to 25 000 from approximately 2.5 million PBMCs (Fig. 2a).

**Figure 2.**
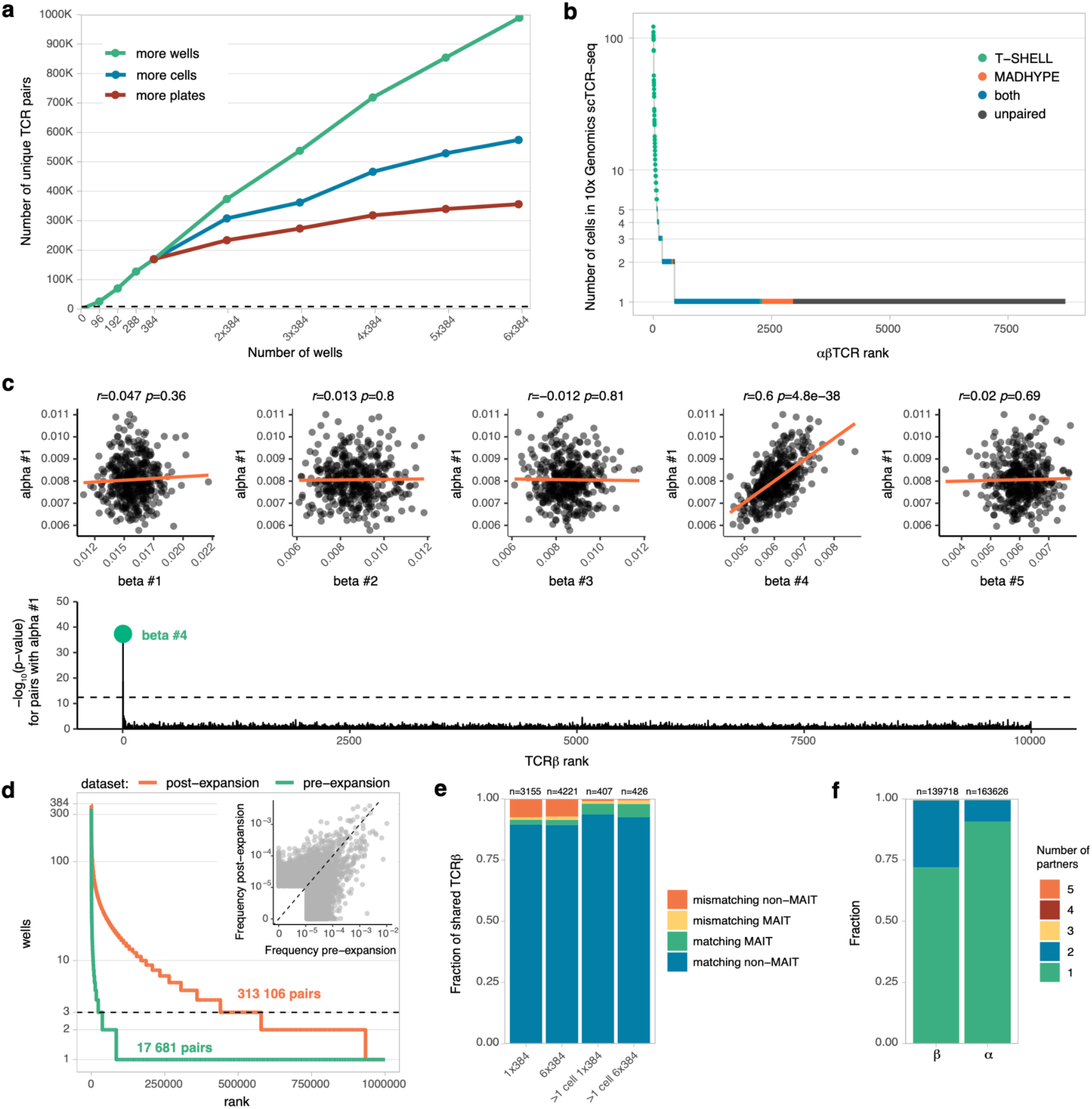
**a.** TIRTL-seq scaling with different analysis methods. The total number of unique paired ɑβTCRs recovered (y-axis) is plotted using three distinct analytical approaches: increasing the number of individual plates analyzed (more plates, red), increasing the number of cells per well analyzed (more cells, blue), or increasing the number of total wells analyzed (more wells, green). **b.** Clone-size distribution of paired and unpaired (dark gray) clones by TIRTL-seq. Paired ɑβTCRs from a 10x Genomics scTCR-seq experiment are ordered by the number of cells with a clonotype (y axis, log scale), while the x-axis shows the clone rank. Each color indicates overlap between 10x Genomics scTCR-seq and ɑβTCRs called by MAD-HYPE (orange), T-SHELL (green) or both (blue). Dark gray indicates lack of pairing in TIRTL-seq compared to 10X Genomics scTCR-seq. **c.** T-SHELL algorithm pairs TCR chain through frequency correlation. **top:** Correlation of relative per well frequencies of largest TCRɑ #1 and top 5 largest TCRβs in the repertoire, red line shows linear fit. **bottom:** Manhattan plot for p-values for pairing of TCRɑ #1 to 10 000 most abundant TCRβs. Dotted line shows p-value cutoff after Bonferroni multiple testing adjustment. **d.** T cell expansion increases pairing efficiency. T cell clone frequency (y-axis) is plotted against rank (x-axis). The dotted line shows the minimal 3-well occurrence threshold for ɑ/β chain pairing by TIRTL-Seq. An increased number of clones clearing the threshold after expansion (orange curve) results in more called pairs compared to pre-expansion (green curve). Inset shows clonal frequency distortion after antigen-independent T cell expansion. Clonal frequency pre-expansion (x-axis) is plotted against clonal frequency post-expansion (y-axis). **e.** Fraction of TCRβs overlapping between 10X Genomics scTCR-seq (filtered or unfiltered for clones with > 1 cell) and TIRTL-seq experiments (x-axis) with matching or mismatching TCRɑ for MAIT and non-MAIT clones. **f.** Fraction of clonotypes with a given chain (ɑ or β) paired with one or more partner chains (TIRTL-seq data from one 384-well plate experiment).

### Extending ɑβTCR pairing to large and small clonal frequencies

To validate TCR chain pairings inferred by TIRTL-seq, we compared results with those from the state-of-the-art scTCR-seq method using the 10x Genomics 5’ GEM-X protocol. We generated 10x Genomics scTCR-seq data for 20 000 T cells from the same donor resulting in 11 113 cells with paired ɑβTCR sequences, corresponding to 8705 unique ɑβTCRs. We compared the paired clonotypes identified by MAD-HYPE analyses of TIRTL-seq data from one 384-well plate with 10x Genomics scTCR-seq to measure the concordance between the two methods. We examined the sizes of clones from 10x Genomics scTCR-seq paired in the TIRTL-seq experiment with the MAD-HYPE algorithm and found that although 2706 ɑβTCRs matched between technologies, no pairs for the abundant T cell clones present at more than 5 cells in the 10x Genomics experiment matched (Fig. 2b, orange and blue dots). Upon further investigation, we found that all these abundant clones were sampled in all or almost all wells of the 384-well plate, thus making it impossible to pair by matching ɑ/β chain occurrence patterns using the MAD-HYPE algorithm. This weakness in the chain occurrence pattern approach was noted in the original MAD-HYPE algorithm and in the pairSEQ protocol (Howie et al. 2015; Holec et al. 2019).

To address this problem, we developed a new heuristic algorithm that we named T-SHELL (**T**CR ɑβ **S**equence **H**ighly **E**fficient **L**inkage **L**earning). The T-SHELL algorithm is based on the fact that clones present in all wells still have variability in the number of cells between wells. Thus the relative frequency of both TCRɑ and TCRβ transcripts within a well should increase if more cells of a given clone are sampled in a given well. T-SHELL employs correlation between TCRɑ and TCRβ clonotype relative frequencies (measured as read fraction) across wells instead of presence/absence patterns to pair them. To demonstrate the principle of T-SHELL, we plotted the correlation between the largest TCRɑ clonotype (by average frequency across wells in the 384-well TIRTL-seq plate) and the top 5 largest TCRβ clonotypes. Read fraction of TCRɑ #1 strongly correlated (Pearson r=0.6, p=10^-38^) with TCRβ #4 (Figure 2c, top scatter plots). TCRβ #4 remained the only TCRβ chain that had a significant correlation with TCRɑ #1 when we correlated the read fractions of TCRɑ #1 against the top 10 000 largest TCRβs (Fig 2c, bottom Manhattan plot). Corresponding 10x Genomics scTCR-seq data confirmed that TCRβ #4 is indeed the correct pair for TCRɑ #1, indicating that T-SHELL can accurately pair large clones. We next applied the T-SHELL algorithm to pair other TCRɑ/β chains found across the 384-well plate and found the majority of abundant clones (73/74 with more than 5 cells per clone on Fig. 2b) are paired and matched the 10x Genomics scTCR-seq pairings (Fig. 2b, blue and green color). Out of 2159 TCRɑβ pairs identified by the T-SHELL algorithm that matched the 10x Genomics scTCR-seq pairs with 1 to 5 cells per clone, 94% (2031/2159) were also identified by MAD-HYPE, suggesting good agreement between the two algorithms (Fig. 2b, blue). Thus, we developed a clonotype frequency-based algorithm capable of pairing TCR chains for abundant clones present in all wells.

For small clones that correspond to the majority of the TCR repertoire in the peripheral blood, TIRTL-seq relies on a probabilistic approach based on occurrence pattern matching. Consequently a clone has to be present in at least 3/384-wells, and thus have at least 3 cells/clone in a sample in order to be paired by TIRTL-seq. To address this limitation and to increase the number of cells per clone, we expanded T cells for one week in the presence of anti-CD3 antibody, IL-2, and IL-15 for donor PBMCs where we had initially identified 17 681 paired clonotypes using the standard TIRTL-Seq protocol, with 8000 PBMCs loaded per well. *In vitro* culturing significantly expanded clones, and consequently increased the number of TCRɑ/β chain pairings by two orders of magnitude from approximately 17 681 to 313 106 pairs (in comparison to the non-expanded sample with the same starting cell count). However, T cell expansion distorted the clonal frequency hierarchy (Fig. 2d): none of the top 5 largest clonotypes pre-expansion were detected within the top 10 post-expansion, and more than ten thousand clones increased or decreased in clonal frequency more than ten-fold.

For further analysis of TIRTL-seq data, we used both MAD-HYPE and T-SHELL algorithms to call pairs and used TCRɑβ pairings identified by either method. For TCRβ clonotypes overlapping between 10x Genomics scTCR-seq and TIRTL-seq, 91.5% (2886/3155) for a single 384-well plate and 91.4% (3858/4221) for the six 384-well plate experiments had the same TCRɑ. Interestingly, many mismatches between TIRTL-seq and 10x Genomics scTCR-seq (15% of mismatches for 6x384 plate experiment) correspond to mucosal-associated invariant T (MAIT) cells, a specialized T cell subset featuring an invariant TCRɑ chain paired with various TCRβs, breaking the TCRɑ/β chain pattern matching in our algorithms (Garner, Klenerman, and Provine 2018). In general, most mismatching ɑβTCRs were found in clones with just one cell in the 10x Genomics scTCR-seq data. The proportion of mismatches dramatically reduced from 8.5% to less than 2% (4/8 mismatches are MAIT) when we selected clones with more than one cell per clone in the 10x Genomics scTCR-seq dataset (Fig. 2e). It has been previously observed that up to 30% of ɑβT cells express two TCRɑ chains (Dupic et al. 2019; Padovan et al. 1993). We identified 27.5% of TCRβ with two TCRɑ partners (Fig. 2f), suggesting TIRTL-seq reliably captures T cells expressing two functional TCRɑ chains.

Thus, the TIRTL-seq approach is as accurate for paired ɑβTCR sequencing as 10x Genomics scTCR-seq at a fraction of the price. TIRTL-seq far exceeds the number of identified TCR pairs compared to the gold standard 10x Genomics scTCR-seq experiment.

### Longitudinal TCR repertoire profiling with TIRTL-seq

Longitudinal TCR repertoire sequencing allows identification of TCR clones expanding and contracting after an immune challenge in an antigen-agnostic manner. To test the performance of TIRTL-seq for longitudinal TCR repertoire profiling, we used three longitudinal samples collected from a SARS-CoV-2-infected individual at -143 days (“baseline”), 6 days (“acute”) and 29 days (“convalescent”) after a positive PCR test (Fig. 3a). On each PBMC sample we performed immunomagnetic positive isolation of CD4+ and CD8+ T cells, which were distributed into two halves of a 384-well plate. We also sorted live CD3+ positive cells from each time point and performed 10x Genomics scRNA-seq and scTCR-seq on ∼20 000 T cells from each time point. Using separate aliquots of PBMCs collected from the same time points, we also performed bulk 5’RACE TCRɑ and TCRβ sequencing (Fig. 3a).

**Figure 3.**
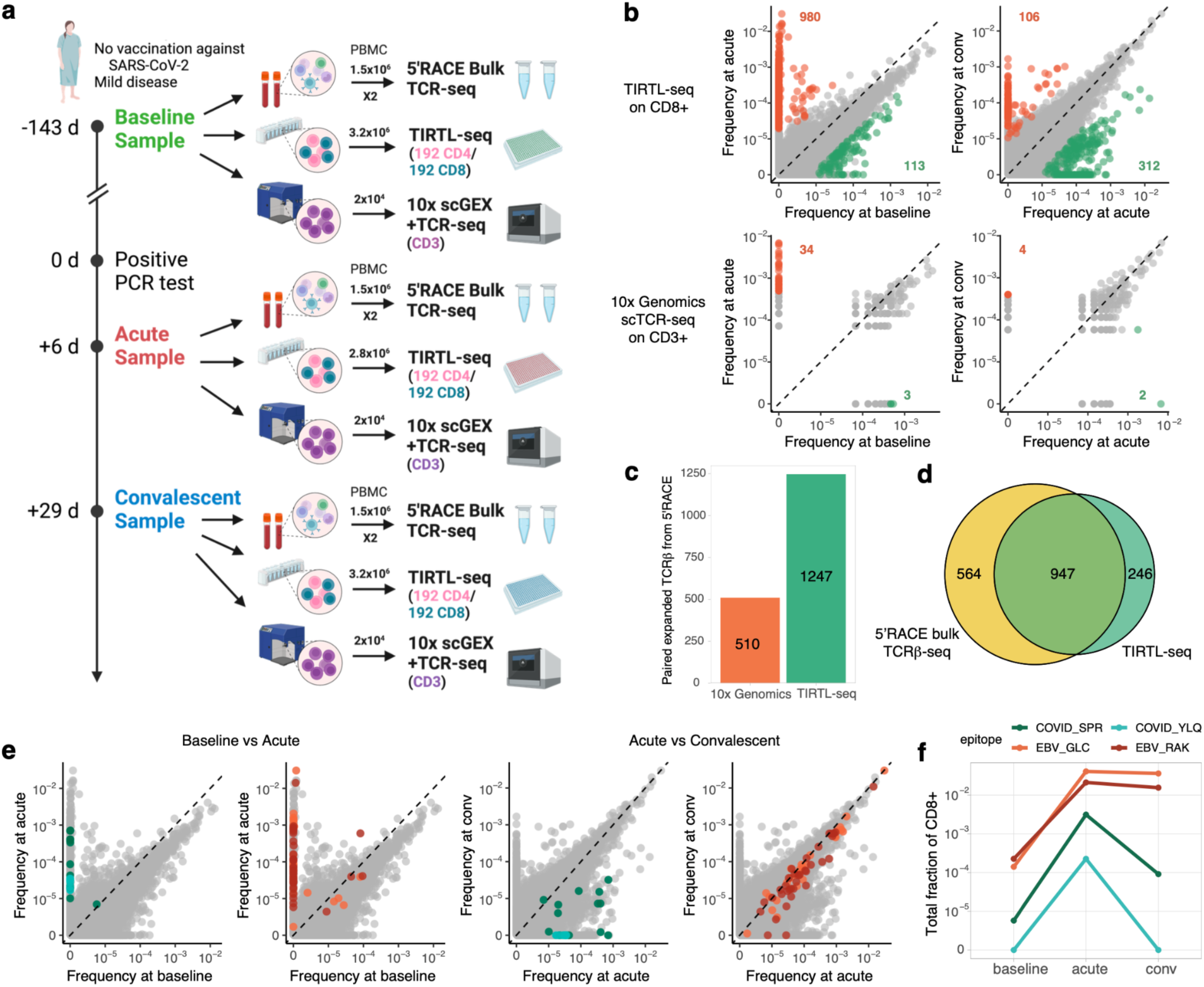
**a.** Longitudinal sampling of a donor with SARS-CoV-2 infection. **b. (top)** TIRTL-seq identifies expansions and contractions in the CD8+ T cell repertoire. **(bottom):** 10x Genomics scTCR-seq clonal frequency from the same time points**. c.** Number of expanded (from baseline to acute) clonotypes from an independent bulk TCRβ sequencing experiment paired using 10x Genomics scTCR-seq and TIRTL-seq**. d.** Overlap between expanded clonotypes from independent bulk TCRβ sequencing (yellow) experiments and TIRTL-seq expanded clones (green, CD4 and CD8 combined)**. e.** Colored dots show clonotypes matching known TCRs specific for A*02 YLQ (cyan) and B*07 SPR COVID epitopes (green) and A*02 GLC (orange) and B*07 RAK EBV epitopes (red) on pairwise time point comparisons. **f.** Cumulative frequency of CD8+ clones specific to A*02 YLQ (cyan) and B*07 SPR COVID epitopes (green) and A*02 GLC (orange) and B*07 RAK EBV epitopes (red) across time points.

To call expanded and contracted clonotypes from TIRTL-seq data, we computed mean frequency and standard error of the mean (SEM) for each TCRβ chain over wells. We call clones significantly expanded or contracted between time points if there is a log_2_ fold change >3 between average frequencies, and the difference between average frequencies exceeds 5 SEM intervals. We identified 980 CD8 (Fig. 3b, top left) and 213 CD4 clones (SI Fig. 1, left) that expanded when comparing the acute to baseline time point. Conversely, 312 CD8 (Fig. 3b, top right) and 134 CD4 clones (SI Fig. 1, right) contracted when comparing acute and convalescent time points. We then applied the same procedure to 10x Genomics scTCR-seq datasets assuming the Poisson distribution for the number of cells with a given TCRβ. While a small number (34) of strongly expanding clones between baseline and acute infection are captured by scTCR-seq (Fig. 3b bottom left), the contraction wave between acute and convalescent is largely undetectable (Fig. 3b bottom right). This is primarily due to the majority of the participating clones having peak frequency of 1:1000, resulting in only a few cells in the 10x Genomics scTCR-seq reaction.

Bulk TCR sequencing is a state-of-the-art method for deep TCR repertoire sequencing and is frequently used in combination with 10x Genomics scTCR-seq to simultaneously identify temporal changes and obtain paired TCR clonotypes. We performed 5’ RACE bulk TCRβ repertoire sequencing, found 1511 expanding clones between baseline and acute replicates, and searched for these expanding clones both in 10x Genomics scTCR-seq and TIRTL-seq results to find TCRɑ partner chains. We found 1247 TCRɑ pairs for expanded TCRβs in TIRTL-seq and 510 pairs in 10x Genomics scTCR-seq (Fig. 3c). The overlap between expanded clonotypes identified by TIRTL-seq and bulk 5’RACE TCRβ-seq was 947/1193 (∼80%), suggesting that TIRTL-seq is efficient both in calling the expansions/contractions and identifying TCR partner chains (Fig. 3d). Importantly, the threshold to call an expanded clonotype between time points is higher than the minimal pairing threshold, leading to the vast majority of expanding clones to also be paired: 887/980 (91%) of CD8 and 195/213 (92%) of CD4 TCRβs increasing from baseline to acute have a paired TCRɑ identified in one of the TIRTL-seq experiments.

Paired ɑβTCR sequencing data generated by TIRTL-seq allowed us to identify some TCRs with known specificities in our dataset. We used TCRdist (Dash et al. 2017) as a pairwise distance metric and found that many clones that expand from baseline to acute time points have highly similar sequences to known TCRs for the HLA-B*07 restricted SPRWYFYYL epitope and HLA-A*02-restricted YLQPRTFLL epitope from SARS-CoV-2 from VDJdb (Goncharov et al. 2022). As expected for SARS-CoV-2 specific T cell responses, these clones contracted from the acute to convalescent time points, indicating death of effector T cells after viral clearance. Interestingly, we also identified 91 expanding clonotypes matching to EBV epitopes (HLA-B*08 restricted RAKFKQLL and HLA-A*02-restricted GLCTLVAML), including the largest clone in the repertoire at acute and convalescent, but not the baseline, time points (Fig. 3e). We cloned two representative TCRs per epitope into NFAT-GFP reporter Jurkat cell lines and confirmed their predicted peptide specificity *in vitro* (SI Fig. 2). In stark contrast to SARS-CoV-2, EBV-specific clones, once established, remained stable from acute to convalescent time points, corresponding to novel clones at the very top of the TCR repertoire (Fig. 3f). We interpret these data to reflect two distinct infections (SARS-CoV-2 and EBV) occurring in this individual during the sample window, as almost none of the epitope-specific responses were detectable at baseline. We measured anti-EBV VCA and anti-SARS-CoV-2 IgG in serum collected alongside the PBMC preparations and confirmed that this donor was seronegative at baseline and seropositive at acute and convalescent time points for both viruses, confirming the longitudinal TIRTL-seq results (SI Fig. 3). These results indicate differential clonal dynamics between acute and chronic viral infections that might explain clonal dominance and the emergence of large stable clones over the lifespan.

## Discussion

Paired TCR sequencing is a powerful approach enabling functional analysis of TCRs and investigation of their interactions with pMHC. Nevertheless, existing approaches for paired TCR sequencing are limited in throughput, expensive, and difficult to implement. Here we present TIRTL-seq, a low-cost methodology for quantitative and deep paired-chain TCR repertoire profiling. Integration of the combinatorial TCR chain pairing strategy suggested by Howie et al. with miniaturization techniques used in the Smart-seq3xpress method (Hagemann-Jensen, Ziegenhain, and Sandberg 2022) allowed us to transition from a 96 to a 384-well plate format and decrease the price per well over 50 times from $2500 per 96-well pairSEQ experiment to $185 per 384-well TIRTL-seq experiment. Additionally, we reimplemented the MAD-HYPE framework (Holec et al. 2019) for GPU, making ɑβTCR pairing inference from experiments with thousands of wells feasible on consumer grade hardware. Furthermore, we developed a new algorithm based on the correlation between the relative abundances of TCR α/β transcripts across wells to overcome the limitations of combinatorial TCR pairing for highly expanded clones sampled in all wells (Howie et al. 2015; Holec et al. 2019; Lee et al. 2017). The result is a method that can process 10 million PBMCs for one-tenth the cost of a standard 10x Genomics experiment, identifying more than 160 000 unique paired ɑβTCRs from a single 384-well plate, albeit without accompanying gene expression data. TIRTL-seq is particularly useful in settings where deep, high-resolution sequencing will be informative, including immune monitoring, tracking responses to immunotherapy, and other immunological interventions including vaccines against infectious diseases and cancer.

Longitudinal tracking of TCR clones can provide biological insight into how the immune system reacts to an immune challenge. For example, here we show different trajectories for SARS-CoV-2- and EBV-reactive TCRs in a set of patient samples before and after confirmed SARS-CoV-2 infection. EBV-reactive clones that were not observed at the baseline time point became the most expanded clones in the repertoire and remained stable at the convalescent time point. While the presence of herpesviruses-specific TCRs among the most abundant clones is consistent with previous reports (Lossius et al. 2014; Huth et al. 2019), here we serendipitously observed how this hierarchy is established in an adult. Further studies into mechanisms that allow certain clones to expand and persist over time will be useful for rational design of vaccine and T cell therapies.

The clonal frequency distribution within TCR repertoires follows a power law. The largest T cell clone in the repertoire typically has a frequency ranging from 1-10%, whereas the frequency drops to about 0.1% for the tenth largest clone, and about 0.01% for the hundredth largest clone (equal to 2 of 20 000 cells loaded into the state-of-the-art 10x Genomics GEM-X reaction). Such low counts make clonal frequency estimation and tracking clones in time challenging with scTCR-seq. TIRTL-seq, on the other hand, allows us to not only determine TCR chain pairings, but also to quantitatively track the frequencies of those clones over time using each well within the plate as a biological replicate. We show that simple averaging of relative clonal frequencies across all wells results in a good clone size estimation. The V-region multiplex RT-PCR strategy without UMIs used in TIRTL-seq is susceptible to amplification bias, however, so incorporating computational adjustments for primer amplification bias could improve the precision of clonal tracking (Carlson et al. 2013).

Our methodology permits chain pairing even for the largest TCR clones in a sample; however, an important limitation is its inability to pair TCRs found in only a few cells. For a 384-well plate experiment, 3 cells per sample is the lowest possible frequency for TCR chain pairing. If pairings of small clones (fewer than 3 cells per sample) are of interest for a particular application, TIRTL-seq can be combined with single-cell sorting to produce paired TCRs for >98% of sorted cells, irrespective of clone size. However, in this scenario the number of analyzed cells is limited by the number of wells in a plate. If sample size is limited to a few thousand cells, conventional scTCR-seq approaches such as 10x Genomics Chromium will likely be more efficient than TIRTL-seq. Alternatively, we show that antigen-independent T cell expansion prior to TIRTL-seq increases the number of cells per clone and thus the pairing probability; this approach, however, disrupts clonal frequency estimates. Low frequency clones are still present in TIRTL-seq results as single TCRɑ and TCRβ chains in large numbers: for 160 000 paired clonotypes in a 384-well TIRTL-seq experiment from 10 million PBMCs we find approximately 5 million and 6 million unpaired TCRɑ and TCRβ, respectively. Unpaired TCR chains could be used to further refine TCR motifs identified in paired results, or as input for any algorithms developed for single chain bulk TCR-seq.

Several methods have been developed to determine TCR specificity from ɑβTCR sequence *in vitro* (Birnbaum et al. 2014; Kula et al. 2019), indicating that all information about TCR specificity is encoded in the paired TCR sequence and suggesting that this problem will eventually be solvable *in silico* (Hudson et al. 2023). Currently, the main limitation for TCR/epitope prediction is the scarcity of training data. Deep, paired TCR repertoire sequencing methodologies are crucial to accumulate data to solve this problem. Once epitope specificity prediction from TCR sequence is solved, affordable and high-throughput paired TCR sequencing techniques will be essential for diagnostics, vaccine, and personalized therapy development. We anticipate that the significant cost reduction and ease of implementation of our method and data analysis pipeline for paired TCR sequencing will have a broad impact and pave the way for its wider adoption, leading to explosive growth in the amount of deep paired TCRɑβ repertoire data. This will contribute both to solving the TCR specificity prediction problem and clinical implementation of TCRɑβ repertoire data for multiple applications.

## Acknowledgments

This work was supported by grants AI136514, AI144616, and AI165077, the St. Jude Center for Influenza Research and Response (SJCEIRR) contract 75N93021C00016, the Center for Influenza Vaccine Research in High Risk Populations (CIVR-HRP) contract 75N93019C00052, the TIRTL Bluesky Initiative, and ALSAC at St. Jude. David C. Brice was supported through The American Association of Immunologists Intersect Fellowship Program for Computational Scientists and Immunologists. The funders had no role in study design, data collection and interpretation, or the decision to submit the work for publication. We thank the Hartwell Center Team at St. Jude, and in particular Scott Olsen, Geoff Neale, and Daniel Darnell for their help with high-throughput sequencing; Greig Lennon at the St. Jude Immunology Flow Core for help with cell sorting; Aleksandra Walczak, Thierry Mora, Phil Bradley, Koshlan Mayer-Blackwell, and Andrew Fiore-Gartland for their valuable discussions of data analysis methods and Dr. Amanda Green, for her advice on serology assays.

## Materials and Methods

### Primary human cells

Healthy donor PBMCs were isolated from apheresis rings obtained from the St. Jude Blood Donor Center under Department of Pathology protocol BDC035. All apheresis rings were de-identified before release. Whole blood was collected from the apheresis ring and adjusted to a total volume of 30 mL with DPBS (Gibco). PBMCs were separated by density gradient centrifugation using Lymphocyte Separation Medium (MP Biomedicals; 10 mL LSM to 30 mL whole blood, 1600rpm at room temperature for 30 minutes, no centrifuge brake). The PBMC layer was then collected, washed with DPBS, and subjected to red blood cell lysis using ACK lysis buffer to remove residual red blood cells. PBMCs were either used immediately for experiments or cryopreserved in Recovery Cell Culture Freezing Medium (Thermo Scientific).

The SJTRC (NCT04362995) is a prospective, longitudinal cohort study involving adult employees (18 years and older) at St. Jude Children’s Research Hospital. The study received approval from the St. Jude Institutional Review Board. All participants provided written informed consent prior to enrollment and regularly completed questionnaires regarding demographics, medical history, treatment, and symptoms if they tested positive for SARS-CoV-2 by PCR. Participants underwent weekly PCR screening for SARS-CoV-2 infection while on the St. Jude campus. Blood samples were collected in 8-mL CPT tubes, processed within 24 hours into cellular and plasma components, aliquoted, and then frozen for future analysis. For our analysis, we selected samples from a healthy donor with naturally acquired mild SARS-CoV-2 infection and no prior history of SARS-CoV-2 infection or vaccination. Samples were collected 143 days before this donor’s first positive SARS-CoV-2 PCR test (“baseline” sample), 6 days after (“acute” sample), and 29 days after (“convalescent” sample).

### Single-cell sorting

Frozen PBMCs were thawed in a water bath and resuspended in 5 mL of prewarmed complete RPMI medium (RPMI 1640 [Gibco] supplemented with 10% FBS [Gibco], 2 mM L-glutamine [Gibco], 100 U/mL penicillin/streptomycin [Gibco]). After centrifugation (500g, 5 minutes, room temperature) the supernatant was discarded, and the cell pellet resuspended in 4 mL of DPBS containing 0.1% BSA and 2mM EDTA. 10 µL aliquot of the cell suspension was used for cell counting with acridine orange and propidium iodide (AO/PI) staining reagent on CellDrop FL (Denovix). Following another centrifugation (500g, 5 minutes) the supernatant was discarded and the cell pellet was resuspended in 50 µL of FACS Buffer (0.5% BSA, 2 mM EDTA in DPBS) with 1 µL of human TruStain Fc-block (1:50, Biolegend) and incubated for 15 minutes at 4 °C. The cells were then stained with 50 µL of a surface antibody cocktail containing Ghost Violet 510 viability dye (1:100, Tonbo Biosciences) and anti-Human CD3-FITC (1:100, Biolegend, clone SK7) for 30 minutes at 4 °C. After washing with DPBS, the cells were filtered through a 50 um filter into sorting tubes with DPBS. Live, single, CD3+ cells were sorted (1 cell/well) using Sony SY3200 into 384-well plates containing cell lysis and RT mix (as described below). Following sorting, the plates were immediately sealed and briefly centrifuged before proceeding with TIRTL-seq.

### T cell expansion

Frozen PBMCs were thawed as described above. After centrifugation (500g, 5 minutes, room temperature), the supernatant was discarded, and the cell pellet resuspended in 5 mL of RP10 medium (RPMI 1640 [Gibco] supplemented with 10% human AB serum [Gemini BioProducts], 2mM L-glutamine [Gibco], and 100 U/mL penicillin/streptomycin [Gibco]). Cells were counted as described above. Following another centrifugation (500g, 5 minutes) the supernatant was discarded and the cell pellet was resuspended to 1 million cells per mL in RP10 medium supplemented with 50 ng/mL anti-human CD3 antibody (Miltenyi, clone OKT3), 3000 IU/mL recombinant human IL-2 (Stemcell Technologies), and 15 ng/mL recombinant human IL-15 (PeproTech). 100 µL cell suspension (100 000 PBMCs) per well was plated in a 96-well round bottom plate. Cells were incubated at 37 °C, 5 % CO_2_ for 7 days.

### CD4/CD8 Selection

Frozen PBMCs were thawed as described above. After centrifugation (500g, 5 minutes, room temperature), the supernatant was discarded and the cell pellet was resuspended in 1 mL Dynabeads Buffer (0.1% BSA, 2 mM EDTA in DPBS). CD8 and CD4 T cells were sequentially positively isolated using Dynabeads CD8 and CD4 Positive Isolation Kits (Thermo Scientific), respectively, according to the manufacturer’s protocol. Briefly, for CD8 selection, 25 µL CD8 Dynabeads were washed with 1 mL Dynabeads Buffer, then resuspended with PBMCs in 1 mL Dynabeads Buffer. PBMCs and beads were incubated with slow continuous rotation for 20 minutes at 4 °C. After incubation, the cell-bead suspension was placed on a magnet for 2 minutes. Supernatant containing the CD8- fraction was then carefully removed and used for CD4 T cell Isolation. The bead-bound CD8+ fraction was washed once with DPBS, then resuspended in 100 µL complete RPMI medium and placed on ice during CD4 selection. For CD4 selection, 25 µL CD4 Dynabeads were washed with 1 mL Dynabeads Buffer, then resuspended with the CD8- supernatant. CD8- cells and beads were incubated with slow continuous rotation for 20 minutes at 4 °C. After incubation, the cell-bead suspension was placed on a magnet for 2 minutes, after which CD4- supernatant was carefully removed. The bead-bound CD4 fraction was washed once with DPBS, then resuspended in 100 µL complete RPMI medium and placed on ice. Both bead-bound CD4 and CD8 populations were then detached from beads by adding 10 µL CD4 or CD8 DETACHaBEAD reagent to the appropriate tube, then incubating with slow continuous rotation for 45 minutes at room temperature. After incubation, tubes were placed on a magnet for 1 minute. Supernatants containing isolated CD4 and CD8 T cells were carefully removed and placed in new tubes, and residual cells were collected by washing the beads with 500 µL complete RPMI twice and collecting supernatant. Isolated CD4 and CD8 T cells were then washed twice with DPBS before proceeding with TIRTL-seq.

### TIRTL-seq Library preparation

Thawed PBMCs were centrifuged (500g, 5 minutes) and the cell pellet was resuspended in 5 mL of DPBS (Gibco). Cells were counted as described above. Following another centrifugation (500g, 5 minutes) the supernatant was discarded, and the cell pellet was resuspended in DPBS (Gibco) to the desired final volume and kept on ice until use.

Reverse transcription and PCR I were performed in 384-well plates pre-loaded with 3 µL of Vapor-Lock (Qiagen). Cell lysis buffer and reverse transcriptase mix (0.4 µL/well) containing 0.1% TritonX-100 (Sigma Aldrich), 0.5 mM dNTP (Thermo Scientific), 0.25 ng/µL Random Hexamer Primer (Thermo Scientific), 2.5 µM Oligo(dT) (Thermo Scientific), 1X Maxima H RT buffer (Thermo Scientific), 5 mM DTT (Thermo Scientific), 1U/µL RNasin Ribonuclease Inhibitor (Promega), and 4 U/µL Maxima H Minus Reverse Transcriptase (Thermo Scientific) was dispensed into 384-well plates followed by cell suspension (0.250 µL/well) using I-DOT Mini (Dispendix). The plate was pulse centrifuged to ensure the lysis buffer containing RT master mix and cell suspension were merged underneath the Vapor-Lock overlay. Reverse transcription was performed by incubating the plate at 42 °C for 5 minutes, 25 °C for 10 minutes, 50 °C for 60 minutes, and 94 °C for 5 minutes.

For multiplex PCR I, an equimolar mix at 1.125 µM each of 88 V-segment forward primers (see SI for complete sequence) containing Illumina Nextera reverse adapter sequence was prepared. For reverse primers, an equimolar mix at 10 µM each of *TRBC* and *TRAC* C-segment specific primers with 6- or 8- nt barcodes for experimental identification and Illumina Nextera forward adapter sequence (see SI for complete sequence) was prepared. Multiplex PCR I master mix was prepared by mixing 0.45 µL of V-segment forward primer mix, 0.2 µL C-segment reverse primer mix with experiment specific barcodes, and 1.25 µL of KAPA2G Fast Multiplex Mix (Roche), and nuclease-free water (Thermo Scientific) to 2 µL final volume. PCR I mix (2 µL/well) was dispensed directly to the RT plate using a MANTIS Liquid Dispenser (Formulatrix) to perform targeted amplification of TCRɑ and TCRβ cDNA. Following the addition of PCR I mix the plate was centrifuged to ensure the PCR mix was merged underneath the Vapor-Lock overlay. The plate was incubated for 3 minutes at 95°C for initial denaturation followed by 20 cycles of 15 seconds at 95 °C, 30 seconds at 59 °C, 1 minute at 72 °C. Final elongation was performed for 5 minutes at 72 °C. Pre- and post-PCR I steps were performed in different rooms to avoid cross-contamination.

For indexing PCR II, 1 µM each of forward and reverse primers containing full length Illumina adapter sequences and 384-well specific unique dual index (UDI) barcodes (see SI for complete sequence) were prepared and stored at -20°C. PCR II mix was prepared by mixing 1 µL of 5X Q5 Reaction Buffer (NEB), 0.1 µL of dNTP Mix (10 mM each, Thermo Scientific), 0.05 µL of Q5 Hot Start High-Fidelity DNA Polymerase (NEB), and nuclease-free water (Thermo Scientific) to 3 µL final volume. PCR II mix (3 µL/well) was transferred in a new 384-well plate containing Vapor-Lock overlay and centrifuged briefly. 1 µL of mix of Illumina UDI primers with well specific barcodes (1 µM each) were stamped into PCR II plate using Viaflo 384 (Integra Biosciences). The PCR I products were ∼1:10 diluted by adding 18 µL of Ultrapure Distilled water using a Welljet dispenser (Integra Biosciences). 1 µL of the diluted PCR I products were transferred into PCR II plate using Viaflo384 (Integra Biosciences). The plate was spun down briefly and incubated at 98 °C for 30 seconds, followed by 15 cycles of 10 seconds at 98 °C, 10 seconds at 58 °C, 50 seconds at 72 °C. Final elongation was performed for 2 minutes at 72 °C. The libraries were pooled in the VBLOK200 reservoir (ClickBio) by spinning the PCR II plate upside down and purified using 1X AMPure XP beads (Beckman Coulter). Final libraries were sequenced on Illumina NovaSeq platform at 300 000 reads per well.

### Manual TIRTL-seq

For the manual version of the protocol, the following changes were made: all steps were carried out in 96-well plates with no Vapor-Lock, all pipetting steps were carried out using standard 8- or 12-well multichannel pipettes, volumes for reverse transcription and PCR I were increased X4 (2.6 µL and 10.6 µL per well final volume respectively), and volumes for PCR II were increased X2 (10 µL per well final volume). Libraries were sequenced on Illumina NovaSeq platform at 2 million reads per well.

### Bulk 5’RACE

Frozen PBMCs were thawed as described above. After centrifugation (500g, 5 minutes, room temperature), the supernatant was discarded and the cell pellet resuspended in 5 mL complete RPMI for counting on Vi-Cell XR automatic cell counter (Beckman Coulter). After counting, the cell suspension was split equally into two replicate aliquots, each containing approximately 5 million viable PBMCs. Following another centrifugation (500g, 5 minutes), supernatant was discarded and the cell pellets were resuspended in 1 mL Trizol (Thermo Scientific) each. RNA was isolated from each replicate sample according to the manufacturer’s protocol and quantified by Qubit RNA High Sensitivity Assay (Thermo Scientific). cDNA synthesis was carried out as described previously (Egorov et al. 2015). Briefly, 250 ng RNA and cDNA synthesis mix containing 1X First Strand Buffer (Takara Bio), 2 mM DTT (Thermo Scientific), 1 mM each dNTP (Thermo Scientific), 1 μM each human TCRɑ and TCRβ RT UMI primers, 1 μM 5’ template-switch adapter, 10 U/µL Smartscribe reverse transcriptase (Takara Bio), and 0.4 U/µL RNasin (Promega) were mixed together, followed by incubation at 42 °C for 60 minutes. After cDNA synthesis, Uracil-DNA glycosylase (NEB) was added at 0.25 U/µL, and samples were incubated for an additional 40 minutes at 37 °C. cDNA was purified using 1X AMPure XP beads (Beckman Coulter).

PCR I reaction mix was prepared by mixing purified cDNA with 1X Q5 polymerase buffer (NEB), 0.2 mM each dNTP (Thermo Scientific), 0.2 μM M1ss forward primer, 0.2 μM each human TCRɑ and human TCRβ UMI first round primers, 0.02 U/µL Q5 Hot Start Polymerase (NEB), and nuclease-free water to a total volume of 50 µL per reaction. PCR I was performed by incubating at 98 °C for 30 seconds, followed by 20 cycles of 98 °C for 10 seconds, 55 °C for 10 seconds, and 72 °C for 50 seconds. Final elongation was performed at 72 °C for 2 minutes. Pre- and post-PCR workflows were performed in separate rooms to prevent cross-contamination.

For PCR II, each sample was split and TCRɑ and TCRβ samples were processed separately. PCR II master mix was prepared by mixing 1X Q5 polymerase buffer (NEB), 0.2 mM each dNTP, 0.02 U/µL Q5 polymerase, and nuclease-free water to a final volume of 50 µL per reaction. Two tubes (one TCRɑ and one TCRβ) were prepared for each sample, and 2 µL unpurified PCR I product, 0.2 μM H1s primer, and 0.2 μM human acj_i (TCR ɑ) or bcj_i (TCR β) was added to each sample. PCR II was performed by incubating at 98 °C for 30 seconds, followed by 16 cycles of 98 °C for 10 seconds, 58 °C for 10 seconds, and 72 °C for 50 seconds. Final elongation was performed at 72 °C for 2 minutes. PCR II product was purified using 0.8X AMPure XP beads (Beckman Coulter).

### 10x Genomics scTCR-seq

PBMCs were stained as described in the **Single-cell sorting** section. Live, single CD3+ cells were sorted on Beckman Coulter Cytoflex SRT sorter, counted on hemocytometer, adjusted to 1400 live cells/uL in DPBS and loaded into 10x Genomics Chromium GEM-X 5’ reaction for target recovery of 20 000 cells. scTCR libraries were prepared by following the manufacturer’s protocol for Chromium GEM-X Single Cell 5’ Reagent Kits v3 (CG000733, Rev A) and sequenced on Illumina NovaSeq platform.

### Raw data processing

Raw fastq files from each well were processed with *mixcr* - v 4.6.0 (Bolotin et al. 2015) package using the *analyze* tool. Optional switches were set to analyze the ‘*generic-amplicon*’ preset with *floating-left-alignment-boundary*, *floating-right-alignment-boundary* for C segment and *split-by-sample*. All other parameters were set to default. *Tag-pattern* switch was used to match and split the samples based on plate barcodes incorporated during the PCR I step. Processing steps included demultiplexing raw reads by plate barcode, aligning V, D and J segments, assembling identical reads into clonotypes and frequency based error-correction.

### Single cell TCR-seq with TIRTL-seq

To call functional TCR chains from single-cell sorted T cells and filter sequencing errors from each well we select functional (no frameshifts and no stop codons inside CDR3) TCR chains with >50 reads and more >10% of reads in a well corresponding to these clonotypes.

### Statistical analysis

#### TCR pairing with MAD-HYPE

Due to variability in cell dispensing and library preparation, some wells produce significantly fewer clonotypes than average. To avoid confounding downstream analysis we exclude wells with less than 50% of the average number of unique TCRɑ clonotypes across the plate.

We reimplemented the MAD-HYPE algorithm (Holec et al. 2019) in R and Python 3. Briefly, for the given experiment design (number of wells and number of cells per well) and TCRɑ and TCRβ overlap statistics (*w_ij_*number of wells where TCRɑ and TCRβ are found together, *w_i_,* number of wells with TCRɑ only and *w_j_,* number of wells with TCRβ only), MAD-HYPE outputs a posterior probability log-ratio for hypothesis that TCRɑ and TCRβ are paired vs. unpaired.

To call TCRɑ/β chain pairs we calculate this ratio for all possible TCRɑ/β pairs and filter all pairs with score above 0.1. To speed up the computations we first compute a look up table containing threshold values for maximal chain loss: for each overlap *w_ij_*, for each loss of TCRβ *w_i_* what is the maximum loss of TCRɑ *w_i_*so paired vs. unpaired hypothesis probability log-ratio is still above the 0.1 threshold.

To efficiently calculate *w_ij_*, *w_i_* and *w_j_* for all possible TCRɑ/β pairs we use GPU and *cupy* (Okuta et al. 2017) or *MLX* frameworks (Hannun et al. 2023), although for the vast majority of 384-well plate TIRTL-seq experiments a single CPU core is enough to analyze the results with our implementation.

#### T-SHELL heuristic algorithm for TCRɑ/TCRβ pairing

If clone is large and is present in all (or almost all) wells, there still should be Poissonian variation in exact number of cell per well, influencing clonal frequency of respective TCRɑ and TCRβ transcripts (fraction of reads corresponding to this clone of all reads) measured in each well.

To pair ɑ/β TCR chains found in all wells, we compute the Pearson correlation coefficient between per well clonal frequencies (defined as sum of reads corresponding to given unique alpha or beta CDR3nt sequence divided by total number of sequencing reads per well) for each possible TCRɑ/TCRβ pair. We next compute the p-value for *H_0_*: *r*=0 (using Student’s t-distribution with *n*-2 degrees of freedom, where *n* is the number of wells). After that for each TCRɑ we order p-values and divide each by third smallest p-value to get adjusted p-value, reflecting our belief that each TCRɑ is paired to at most two TCRβ sequences. We then put a 10^-10^ threshold on the adjusted p-value to call chain pairs. We previously suggested a similar approach for alpha/beta pairing by longitudinal clonal frequency trajectories matching (Minervina et al. 2020).

#### Identification of expanded/contracted clones

To identify expanded/contracted clones between timepoints with TIRTL-seq we first average relative frequency (defined as sum of reads mapping to given CDR3β divided by sum of reads mapping to all CDR3βs) across all wells in a plate, including wells were the clone is not present. We filter out clones found in less than 5 wells in either time point to limit the number of comparisons. For each clonotype we calculated standard error of the mean on both timepoints, and called clones significantly expanding/contracting if log_2_ fold change between average frequencies on two timepoints is larger than 3 and 2.5 SEM intervals do not overlap. If clone has very low frequency on one of the timepoints and found in fewer than 3 wells we assumed that SEM is twice SEM of clones found in 3 wells at this time point, and pseudocount of 10^-6^ was added to all clonal frequencies to get reasonable fold change estimates.

To identify expanded/contracted clones from bulk TCRβ repertoire replicates we used *edgeR* package (Robinson, McCarthy, and Smyth 2010), as described before (Pogorelyy et al. 2018), using p.adj<0.001 and log2FC<2 thresholds.

#### TCR specificity matching to VDJdb

To identify clusters of highly similar clones matching previously reported TCR specificities, we co-cluster paired data from acute timepoint with VDJdb and identified tightly interconnected clusters including both TCRs with previously described specificity and abTCRs from our experiment. First we filtered VDJdb (accessed on Aug 6 2024) for paired human TCRs with reported HLA-restriction matching HLA class I alleles of our donor (A*02, A*03, B*07, B*08 and C*07). We excluded A*02 GILGFVFTL and A*03-restricted KLGGALQAK epitopes to avoid spurious matching. We then merged the cleaned-up VDJdb dataset with paired TCRs detected on acute time points and used TCRdist<90 to connect similar clones with edges. To separate tightly interconnected clusters of highly similar TCRs we excluded 1% of nodes with highest betweenness values from the network. We then assign specificity to TCR clusters if more than 40% of nodes from the cluster were from given epitope-specific TCRs from VDJdb.

#### TCR cloning and screening

TCR cloning and screening was performed as previously described (Minervina et al. 2022; Mudd et al. 2022). Briefly, to generate artificial antigen-presenting cells, gBlock gene fragments encoding HLA-A*02:01, HLA-B*08:01, and HLA-B*07:02 were synthesized by GenScript and cloned into a pLVX-EF1α-IRES-Puro lentiviral vector (Clontech). Lentivirus was produced by transfecting HEK 293T cells (ATCC CRL-3216) with the HLA-containing vector, psPAX2, and pMD2.G plasmids (Addgene) at a ratio of 4:3:1 using Lipofectamine 3000 kit (Thermo Fisher Scientific). Viral supernatant was filtered, concentrated, and used to transduce K562 cells (ATCC CCL-243), followed by selection with puromycin for 1 week. Surface HLA expression was confirmed by flow cytometry.

Representative TCRɑβ pairs for each epitope were selected (see SI Table 2), modified with murine constant regions, and linked to mCherry via 2A sites. This construct was cloned into a pLVX-EF1α-IRES-Puro vector. Lentivirus was generated by transfecting HEK 293T cells (ATCC CRL-3216) with the TCR-mCherry vector, psPAX2, and pMD2.G at a ratio of 4:3:1 using Lipofectamine 3000 kit (Thermo Fisher Scientific). 2D3 Jurkat 76.7 cells expressing human CD8+ and NFAT-GFP (Morimoto et al. 2018) (a kind gift from Fumihiro Fujiki), were transduced and selected with puromycin for 1 week.

Resulting TCR-transgenic Jurkat cells (10^5^) were co-cultured with 10^5^ K562 aAPC cells with a single predicted HLA in a round-bottom 96-well cell culture plate in 100 µL RPMI-1640 media (Gibco) supplemented with 10% FBS, 1% penicillin-streptomycin, and 1% L-glutamine, and pulsed with predicted peptide at 10 µM final concentration. Predicted HLA-B*07:02-SPRWYFYYL-specific TCRs were additionally tested against LPRWYFYYL, an epitope variant found in HKU1 and OC43 common cold coronaviruses. Fraction of GFP+ cells of mCherry positive cells was measured on BD FACSymphony A3 flow cytometer.

#### anti-EBV and anti-SARS-CoV-2 ELISA

Anti-SARS-CoV-2 IgG analysis was performed as previously described (Lin et al. 2022). Briefly, 384-well microtiter plates were coated overnight at 4 °C, with recombinant SARS-CoV-2 Spike (Sino Biological) diluted in PBS at 2 µg/mL. Plates were washed three times the next day with PBS-T (0.1% Tween-20) before being blocked with 3% Omniblok non-fat milk (AmericanBio) in PBS-T for one hour. Plates were washed as before, then incubated with the serum samples diluted 1:50 in 1% milk in PBS-T for 90 minutes at room temperature. Prior to dilution, serum samples were thawed and heat-inactivated at 56 °C for 15 minutes. The plates were washed as before and incubated for 30 minutes at room temperature with anti-human IgG secondary antibody (Invitrogen) at a 1:10,000 dilution in 1% milk in PBS-T. After plates were washed as before, they were incubated with SIGMAFAST OPD (Sigma-Aldrich) for eight minutes in the dark at room temperature. To stop the chemiluminescence reaction, 3N HCl was added to the wells of the plate. The plates were then read at 490 nm on a microplate reader. Binding antibody units per mL (BAU/mL) were determined by comparing the ODs of the target samples to those of blank wells and samples calibrated to WHO standard samples (NIBSC) set at 1000 BAU/mL. A cutoff value of 90.9 BAU/ml was used to determine anti-SARS-CoV-2 Spike IgG positivity. This value has been used by others when measuring these antibody levels utilizing the same WHO control (Winichakoon et al. 2023) and was slightly higher than the values of other baseline serum samples from SJTRC participants. For anti-EBV VCA IgG analysis, serum samples were prepared and measured according to the manufacturer’s instructions (Abcam ab108730). Standard units (SU) were calculated using the provided control samples measured OD at 450 nm. The ODs of the control samples and blanks were within the manufacturer’s criteria for a valid assay run.

### Data and code availability

TCR sequencing data is available at Zenodo (**10.5281/zenodo.14010377**), code is available at GitHub (https://github.com/pogorely/TIRTL).

## Supplementary Information

**SI Fig. 1.**
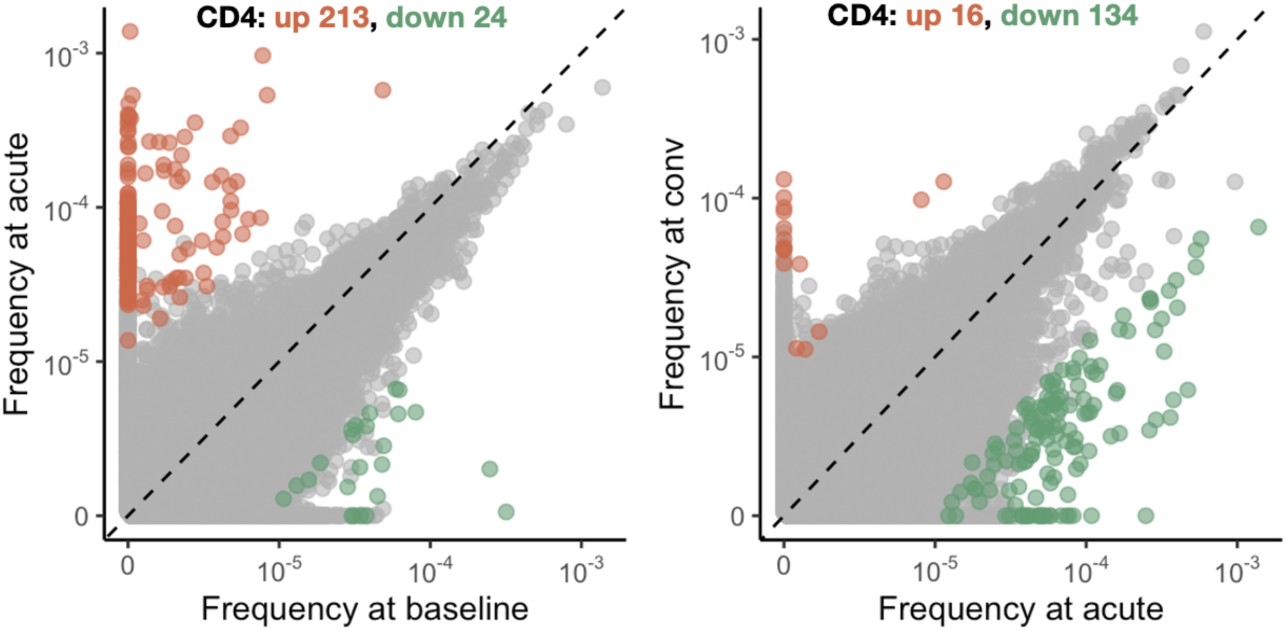
TIRTL-seq identifies expansions and contractions in the CD4+ T cell repertoire. Each dot represents TCRβ clonotype, frequency at two timepoints is plotted in log-scale. Orange and green color show significantly expanding and contracting clones respectively.

**SI Fig. 2.**
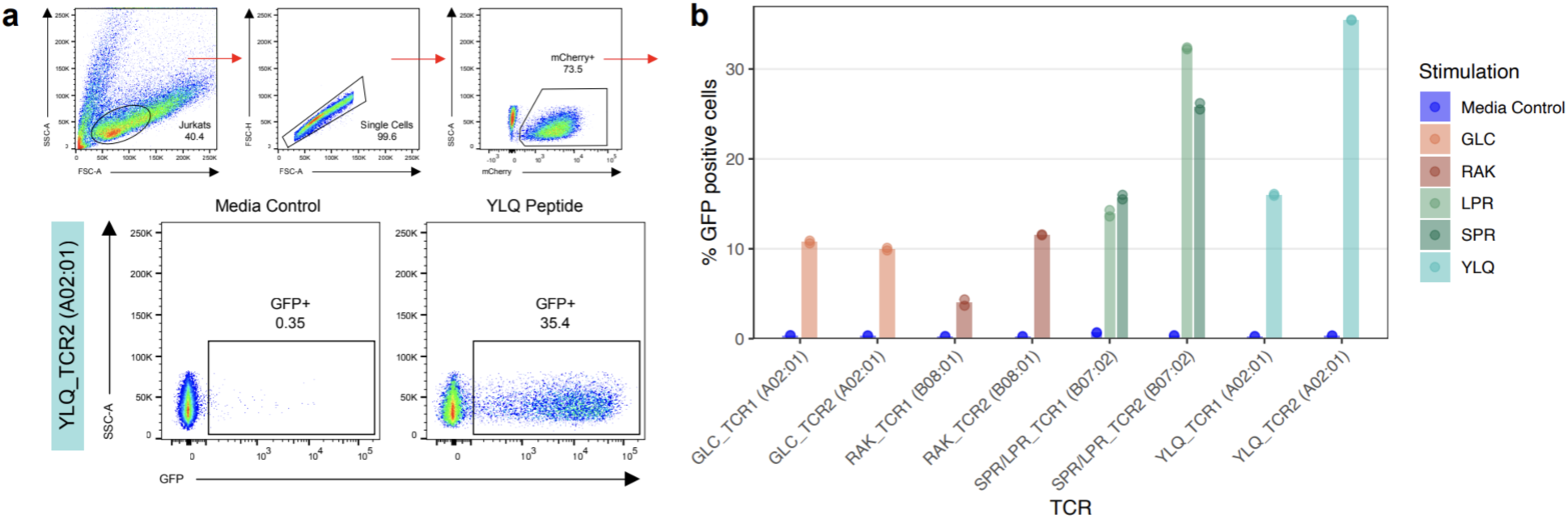
**a.** Gating strategy and representative flow plots for *in vitro* validation experiments. **b.** *In vitro* validation of predicted TCR specificity. Bars show average percentage of GFP+ cells out of TCR-transgenic Jurkat cells (mCherry+), dots show replicates.

**SI Fig. 3.**
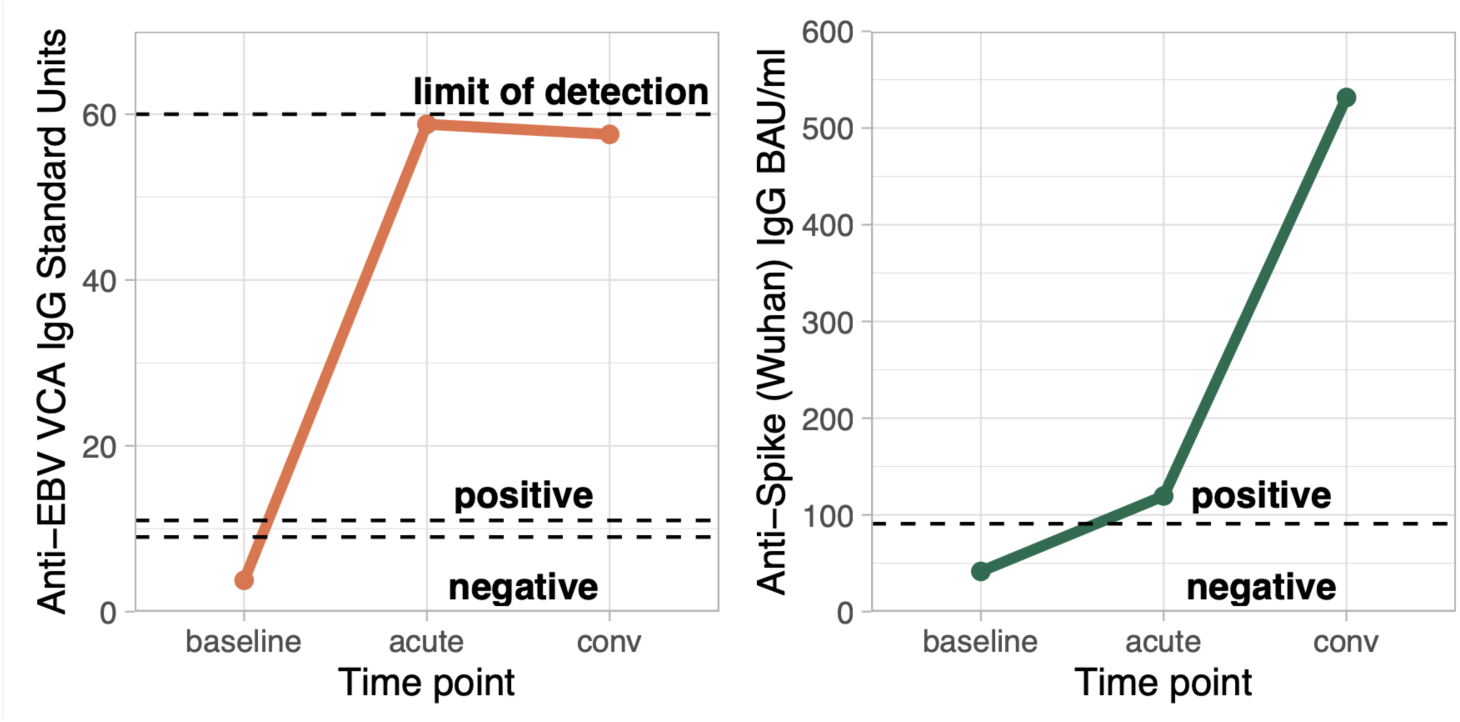
Anti-EBV VCA (left) and Anti-SARS-CoV-2 Spike IgG levels. Lowest dashed line shows seronegativity cut-offs.

